# L-lactic and 2-ketoglutaric acids, odors from human skin, govern attraction and landing in host-seeking female *Aedes aegypti* mosquitoes

**DOI:** 10.1101/2022.10.18.512748

**Authors:** Benjamin D. Sumner, Brogan A. Amos, Jan E. Bello, Ring T. Cardé

## Abstract

*Aedes aegypti*, presented with a source of L-lactic and 2-ketoglutaric acid in a wind-tunnel bioassay, takeoff, fly upwind, and land on the blend at rates comparable those exhibited by mosquitoes presented with a skin odor stimulus. Addition of carbon dioxide decreased takeoff latency but was not required to elicit upwind flight nor landings. Ketoglutaric acid, a recently identified component of human skin odor, combined with lactic acid elicits the full repertoire of mosquito host-seeking behaviors.

## Introduction

*Aedes aegypti* (Diptera: Culicidae) is a vector of several consequential arboviruses including chikungunya, dengue, yellow fever, and Zika. *Aedes aegypti* has invaded much of the tropics and sub-tropics, making dengue the most prevalent human arbovirus, infecting 100 million per year and placing nearly half of the world’s population at risk (Bhatt et al. 2013).

Females of the anthropophilic form of *Ae. aegypti* blood feed almost exclusively on humans (Scott et al. 1993). This makes them particularly effective vectors of human pathogens (MacDonald 1952). The cosmopolitan “subspecies” *Ae. aegypti aegypti* diverged from the non-human preferring *Ae. aegypti formosus*, 400 to 550 years ago (Gloria-Soria et al. 2016; Crawford et al. 2017; Powell et al. 2018). Hereafter references to *Ae. aegypti* are to the anthropophilic Orlando strain of *Ae. a. aegypti*.

To find a host, *Ae. aegypti* fly upwind when they detect fluctuating levels of CO_2_ above ambient concentration. Once they are within several meters of a prospective host, they can also sense visual cues and, even closer to the host, thermal cues (Gillies 1980; Cardé and Gibson 2010; van Breugel et al. 2015; Cardé 2015; Sumner and Cardé 2022). The precise distances at which they detect and use host odors other than CO_2_ is unresolved (Gillies and Wilkes 1970; Dekker et al. 2005).

For *Ae. aegypti*, humans are distinguished from other potential hosts by their skin odor (Steib et al. 2001; Dekker et al. 2002; McBride et al. 2014). Bernier et al. (2002) characterized 279 compounds in the headspace above human skin. A review by Dormont et al. (2021) provides a valuable table of mosquito attractants using Dethier et al.’s (1960) definition of attractant as “a chemical which causes insects to make oriented movements towards its source.” Their list of compounds is organized by species, “co-tested” compounds, and assay type. A challenge is to determine which compounds or combinations of compounds elicit host-finding behaviors. This task is complicated by how different bioassays measure different components of mosquito host-seeking behavior. “Attractiveness” in one assay may measure flying into a port from a still-air chamber, whereas in another, it may measure arrestment after upwind orientation (Cardé 2022). Most assays measure attraction by the responder reaching an endpoint of orientation over a set interval.

Intermediate steps in orientation also can be monitored. Dekker et al. (2001) in a still-air, port -entry assay and Torr et al. (2008) in a field trapping study used electrocution grids to show that some host odors can lure mosquitoes to the vicinity of the odor source but do not always elicit port or trap entry, respectively. The reasons for such outcomes may relate to the odors being incomplete or containing some antagonistic compounds. Alternatively, the odor plume’s structure may be suboptimal for orientation (Geier et al. 1999; Dekker et al. 2001). Another metric to monitor is the rapidity of orientation. Dekker et al. (2001) documented the time to either electrocution or port entry, showing that some odor blends induced more rapid orientation than others, but if the assay’s duration was extended to 15 min, this difference faded. Kennedy (1977) discussed the importance of the assay’s duration in its outcome and the limitation of endpoint assays in distinguishing undirected movement (kinesis) from directed movement (taxis).

Nonetheless, laboratory endpoint assays are useful for investigations of mosquito responses to host emitted odors. For example, BG-Lure, a human skin-odor mimic, designed for use with the Biogents Sentinel® trap (Biogents, Regensburg, Germany), releases lactic acid, ammonia, and hexanoic acid. This lure was developed using a Y-tube olfactometer with subsequent field-trapping studies (Williams et al. 2006).

Lactic acid (2-hydroxypropanoic acid) is thought to be a diagnostic cue used by *Ae. aegypti* (Steib et al. 2001) and *An. gambiae* (Dekker et al. 2002) to distinguish humans from non-human animals. The addition of lactic acid made non-human animal odor more attractive to both species of anthropophilic mosquitoes. Both studies found lactic acid, however, was necessary but insufficient by itself to attract the number of *Ae. aegypti* that were attracted to human odor. Smith et al. (1970) tested *Ae. aegypti* in a combined port entry and landing assay. They found that lactic acid reduced landing on a human-worn sock, but increased olfactometer trap catch downwind of the sock, suggesting that the dose of lactic acid reaching the mosquito is important. Healy and Copland (2000) found that *An. gambiae* did not land on a lactic acid source. However, Dekker et al.’s (2002) findings, lactic acid may be an attractant but not a landing cue for *An. gambiae*.

Recently, Bello and Cardé (2022) identified 2-ketoglutaric acid (2-oxopentanedioic acid) in human skin odor and demonstrated that a mixture of lactic and ketoglutaric acids in the presence of CO_2_ is a landing cue for *Ae. aegypti*. The addition of pyruvic acid, also present in the active fraction of skin odor (Bello and Cardé 2022), to this mixture did not significantly elevate landing rates above those elicited by a blend of lactic and ketoglutaric acids.

Carbon dioxide added to ambient air (which has an intrinsic concentration of CO_2_ of about 0.4%), increases the takeoff rate of *Ae. aegypti* (Daykin et al. 1965), elicits upwind flight (Kennedy 1940), increases heat-seeking (McMeniman et al. 2014), and ultimately leads to increased endpoint capture (Huffaker and Back 1943. In a generally accepted model of mosquito host seeking, formalized by Gillies in 1980, CO_2_ is generally considered as the long-range attractant of host-seeking mosquitoes. The model assumes a mosquito detects CO_2_, takes off, flies upwind along the plume, and then switches to other host cues. Dekker et al. (2005) found that diluted human skin odor was less attractive to *Ae. aegypti* than undiluted skin odor and that CO_2_ sensitizes mosquitoes to other host odors. These models were updated by Cardé and Gibson (2010), van Breugel et al. (2015), and Cardé (2015), but still posit that CO_2_ was the long-distance attractant, and that as mosquitoes fly closer to their hosts they would switch to using specific cues such as skin odor.

This paradigm may not hold for other mosquitoes. Schreck et al. (1972) found that *Anopheles quadriannulatus* flew to a calf from further distances than to a calf-equivalent quantity of CO_2_. It is unknown how far downwind the lactic and ketoglutaric acids are detectable by *Ae. aegypti*. Additionally, it is not known whether there are characteristic behaviors, such as surging upwind or casting crosswind, associated with the detection and subsequent attraction to the landing cue compounds

As the blend of lactic and ketoglutaric acids elicited landing in a cage assay (Bello and Cardé 2022), we set out to determine if the blend elicits upwind flight in a wind tunnel in the presence or absence of CO_2_. We used wind-tunnel assays with videography allow examination of orientation maneuvers, in addition to landing (Lacey and Cardé 2011). Some wind-tunnel assays have used video tracking without landing counts (van Breugel et al. 2015); however, landing is the ultimate measure of successful orientation to a host. If a mosquito does not land, it cannot bite nor transmit pathogens (Reed et al. 1900). We measured time from release to takeoff, time from takeoff to the first landing on the odor source, the number of landings, and the duration of landings. Our video tracking system allowed us to examine mosquito flight maneuvers prior to landing.

## Materials and Methods

### Mosquito Rearing

An Orlando strain colony of *Ae. aegypti* was maintained in a L:D 14:10 h photocycle at 27 °C and 70 % RH. Approximately 50 larvae were reared in plastic containers (26 × 25.6 × 15 cm) with ~1 cm of deionized (D.I.) water and fed Tetramin® pellets (Tetra, Blacksburg, VA, U.S.A). Pupae were held in plastic containers, transferred to screen cages (30 × 30 × 30 cm, BugDorm-1, Megaview Science Co. Ltd. Talchung, Taiwan) before eclosion. Mosquitoes were provided 10 % sucrose solution in D.I. water on a cotton wick. Males and females were held together in the screen cages, and females used in the bioassays were assumed to have mated. Females used in experiments were 3-10 days post-eclosion and were not blood fed. Mosquitoes were starved and deprived of water approximately 12 hours prior to experiments. Female mosquitoes were transferred individually to clean cylindrical acrylic release cages (7 × 8 cm diameter) 30 minutes prior to testing; assays were conducted 4-8 h into their photophase.

### Assay Methods

The assay methods were adapted from Sumner and Cardé (2022). The flight and landing of mosquitoes were released in a glass wind tunnel 122 × 30.5 × 30.5 cm and were video recorded (FDR-AX53, Sony, Tokyo, Japan) from above. Air was drawn into the tunnel from an adjacent, uninhabited room (25 °C and 70 % RH). To simulate the presence of an upwind vertebrate host, 100 ml/minute of 4 % CO_2_ mixed with tank air (equivalent to 1/60 of the exhalation a human, Snow 1970), was carried to the wind tunnel via a 3-m-long Tygon® tube, ensuring temperature equilibration (Pinto et al. 2001). The tube was connected to an L-shaped glass tube (OD 5.5 mm, ID 3.5 mm) that descended 15 cm from the ceiling of the tunnel and extended 20 cm downwind to 60 cm upwind from the release cage. The 4 % CO_2_ mix exited at ~0.4 m/s but produced no detectable difference in wind speed (Omega HHF 52 anemometer, Omega Engineering, Inc., Stamford, CT, USA) nor a temperature difference (to within 0.1 °C) 1 cm downwind of the release point. The CO_2_ release tube was centered so that the generated plume of CO_2_ passed over the beads treated with skin odor and then to the release cage. In trials without the addition of CO_2_, tank air was supplied at the same rate through the same equipment.

The assay room was maintained at 27 °C and 60 % RH. Illumination for videography was provided by infrared LED lights (AXIS T90A, 850 nm, Axis Communications AB, Lund, Sweden) mounted behind a stainless-steel screen at the downwind end of the tunnel as well as beside the wind tunnel. The infrared light in the camera was turned off to avoid glare. Visible light was provided by incandescent bulbs and measured at ~14 lux inside the tunnel.

Treatments were presented on glass beads (black, 10/0 Czech Glass Seed, approximately 2 mm OD toroidal, Precosia Ornela, Zásada, Czech Republic) placed in a clean glass Petri dish (7 cm diameter). Negative control beads are hereafter called clean beads. The blend components, ketoglutaric acid (“KGA” in the figures) (0.5 ml; 10 μl/ml or 100μl/ml in acetone) and lactic acid (“LA” in the figures) (0.5 ml; 10 μl/ml or 100μl/ml in acetone) were applied in a dropwise spiral to beads. As Ghaninia et al. (2019) found that acetone was attractive in a flight tube to *Ae. aegypti*, the beads were placed under a fume hood for 10 minutes to ensure that the acetone had evaporated.

Human skin odor was collected onto glass beads by placing 25 ml of beads into a polyester/cotton blend sock, which was worn by BDS for 12 hours. Beads were cleaned after use by soaking in a solution of 10 % detergent (Micro 90 Cleaning Solution, Cole-Parmer, Vernon Hills, IL, USA) in D.I. water and sonicated for one hour. The beads were then thoroughly rinsed with D.I. water, dried, rinsed twice with acetone (ACS grade, Fisher Scientific, Pittsburg, PA, USA), and heated to 250 °C for 12 hours before reuse.

Differing from Sumner and Cardé (2022), the dish of beads was presented on a 15-cm high metal stand in the center of the tunnel, 55 cm upwind of the release cage. This ensured that compounds emanating from the beads were detectable by the mosquitoes in their release cage. Assays were run and recorded with video for 6 minutes, commencing with the opening of the release cage. Disposable nitrile gloves were always worn by the experimenter to prevent contamination with skin odors.

### Treatment Strategy

We presented the mosquitoes with clean beads, a low dose blend of 5 μg each of ketoglutaric acid (KGA) (2-Keto-glutaric acid 97 %, TCI, Tokyo, Japan), and lactic acid (LA) (L-lactic acid 85-90 % in water, Honeywell Fluka, Charlotte, NC, USA) (based on Bello and Cardé 2022), a blend of 50 μg each of lactic and ketoglutaric acids, and skin odor-treated beads by being worn in a sock. All four treatments were tested in the presence and absence of a turbulent 4 % plume of CO_2_ (Table 1).

**Table 1.** Treatment Combinations for Wind Tunnel Assays with Female *Ae. aegypti*.

| **Treatment** | **CO<sub>2</sub>** | **Assays** | **Assays with takeoff** | **Mosquitoes per assay** | **Total mosquitoes** |
| --- | --- | --- | --- | --- | --- |
| Clean beads | No | 46 | 87% | 1 | 46 |
| Clean beads | Yes | 32 | 88% | 1 | 32 |
| 5 µg each of LA+KGA | No | 40 | 75% | 1 | 40 |
| 5 µg each of LA+KGA | Yes | 39 | 90% | 1 | 39 |
| 50 µg each of LA+KGA | No | 32 | 88% | 1 | 32 |
| 50 µg each of LA+KGA | Yes | 39 | 100% | 1 | 39 |
| Skin odor | No | 31 | 74% | 1 | 31 |
| Skin odor | Yes | 29 | 83% | 1 | 29 |
| 50 µg each of LA+KGA | Yes | 12 | All had $\geq 1$ | 5 | 60 |
| 50 µg LA only | Yes | 12 | All had $\geq 1$ | 5 | 60 |

To confirm the blend was not eliciting mosquito landing solely due to its lactic acid content, we also tested 50 μg of LA alone and the blend of 50 μg each of both compounds in a series of one-choice assays with CO_2_. Five mosquitoes were used in each replicate of these assays (Table 1), which enabled a five-fold reduction in the number of assays for these treatments. The potential number of landings was not lowered but criteria such as time to takeoff and time from takeoff to first landing had their sample sizes reduced five-fold.

### Data Acquisition

Video files were observed, and behavior was scored with BORIS v.5.1.0 (Friard and Gamba 2016). All videos were viewed and scored from the release time until 6 minutes elapsed. The observer recorded release and takeoff as “point” events, whereas landing was scored as a “state” event starting with the landing on the beads and ending with the takeoff from the beads.

Video flight tracking was performed using EthoVision XT v.9.0 (Noldus Information Technology, Wageningen, The Netherlands). Raw numeric data were exported and used in statistical analysis. For data obtained with EthoVision XT, tracking commenced at takeoff and continued until the individual landed on the beads or remained in the upwind section of the wind tunnel, and was therefore indistinguishable, for ≥ 30 seconds.

### Statistical Analysis

All data manipulation and statistical tests were conducted using R v.3.5.0 (R Core Team 2013) in RStudio v.1.1.463 (RStudio Team 2020).

#### Landing Observations

The proportion of trials with at least one landing were compared across all treatments with a Fisher Exact test followed by pairwise Fisher Exact tests with Benjamini-Hochberg correction to reduce the false discovery rate. This method considers all the treatments to be completely independent of each other.

All other tests of manual landing observation data were conducted with generalized linear models (GLMs). A matrix was manually constructed with the independent variable data of each of the treatment combinations. It contained binary values for skin odor and CO_2_ as well as a values of 0, 1, or 10 for the dose of the lactic and ketoglutaric acid blend. Instead of considering all treatments as independent, as the Fisher Exact test does, the GLMs used this information about the relationships among the treatments. In particular, the models treat the different doses of the blend as different values of the same independent variable. The GLMs show which treatments were significantly correlated with whichever behavioral outcome was tested. This allowed us to determine which treatments were correlated. If we had relied on testing differences among treatment combinations, we would have potentially masked the importance of some cues. Significant correlations, unlike significant differences, do not translate directly to graphs of whole data sets. Therefore, instead of visually clear asterisks, the outputs of the GLMs are solely listed in the figure captions.

First, a GLM was used to compare the number of trials with at least one landing. This test of the same data examined with the Fisher tests allows comparison across methods. The treatment matrix was used again when comparing the: number of mosquito landings on beads per trial, durations a mosquito remained after landing among the different treatments, latency (time from release to takeoff), and duration of flight from takeoff to first landing. For testing of repeated landings, trials with one landing were converted to arbitrarily small values. The data was root ten transformed to make the residuals acceptably close to normal.

#### Flight Tracks

Kruskal-Wallis tests were used to contrast the mean distance of the mosquito from the beads every 1/15 of a second during flight track, the mean velocity of the mosquito during flight, the E_max_ (track straightness) of the mosquito flight, and the proportion of time the mosquito spent heading (± 20 °) towards the center of the beads. Spearman rank correlations were used to test effects.

E_max_ (a measure of straightness, 1 being a completely straight track) (trajr package, McLean and Skowron-Volponi 2018) was used as a measure of track sinuosity and was calculated (Cheung et al. 2007) using the X and Y coordinates of the subject at each time point. E_max_ is bases on an iterated summing of the expected displacement. As the shortest path between two points is a straight line, a high E_max_ represents a straight path. This displacement does not necessarily correlate to distance from the beads. A mosquito flying a figure-eight over the beads would have a high E_max_ and a low mean distance to the beads before landing, whereas a mosquito that flew straight to the beads would have a low E_max_ and a low mean distance to the beads.

## Results

### Takeoff

The presence of CO_2_ was the only treatment component that was positively correlated with the proportion of trials with mosquito takeoff (Est. = 0.447, P = 0.022) (Fig. 1).

**Fig. 1.**
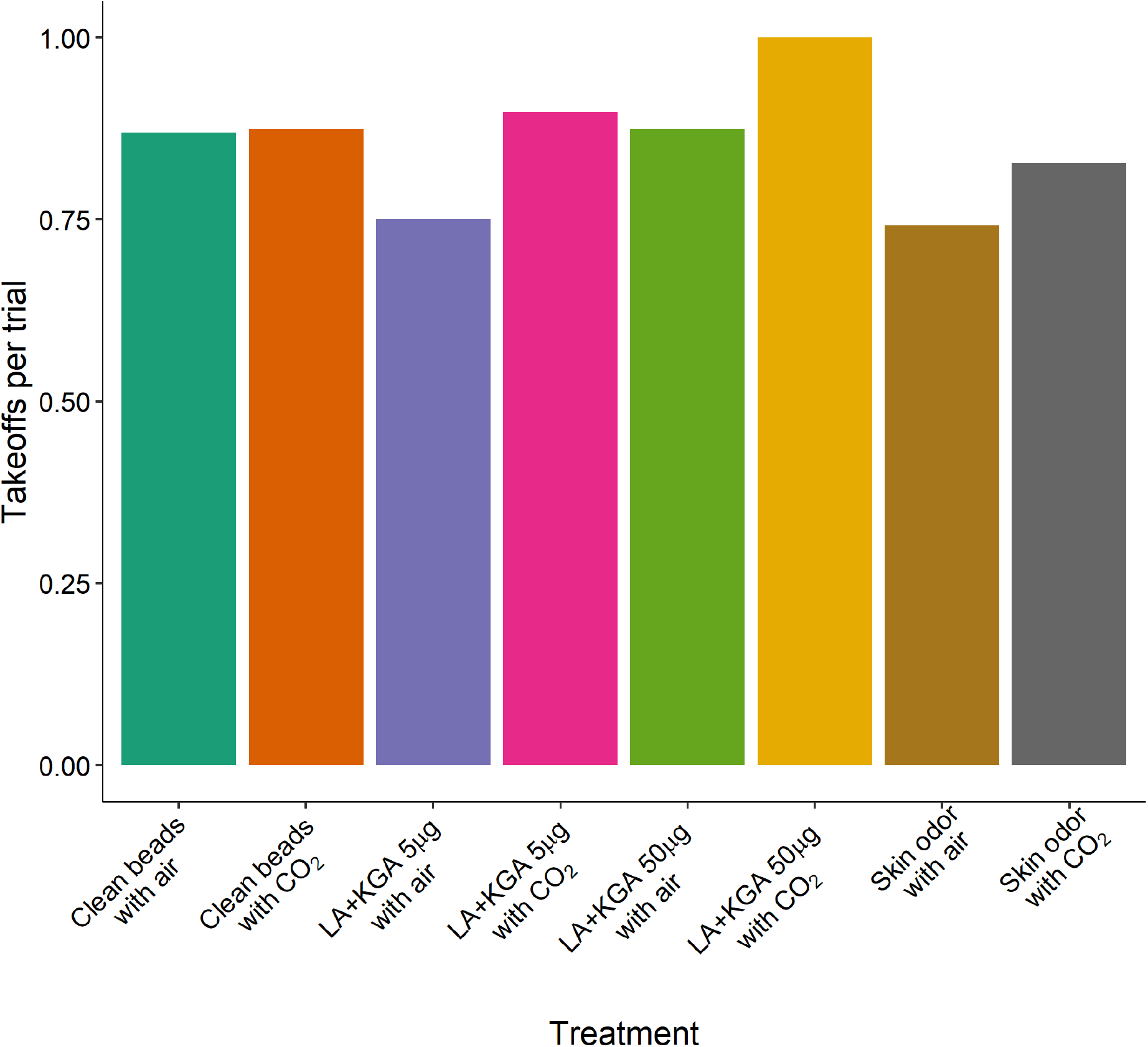
The proportions of trials in which *Aedes aegypti* initiated flight. Only the presence of CO_2_ was correlated with the proportion of trials with takeoff (GLM, binomial, link = probit, Est. = 0.447, P = 0.022).

### Latency from Release to Takeoff

Among the treatments there were no significant differences in takeoff latencies. Although the 50 μg dose appears to elicit a significant decrease in take-off latency (P = 0.017, Coefficient = −0.3534), the residuals of the GLM (Gamma, link = log) were non-normal (KS P = 0.022), which means that the test cannot be used in this case (Fig. 2). This distribution and link function produced the closest to normal residuals.

**Fig. 2.**
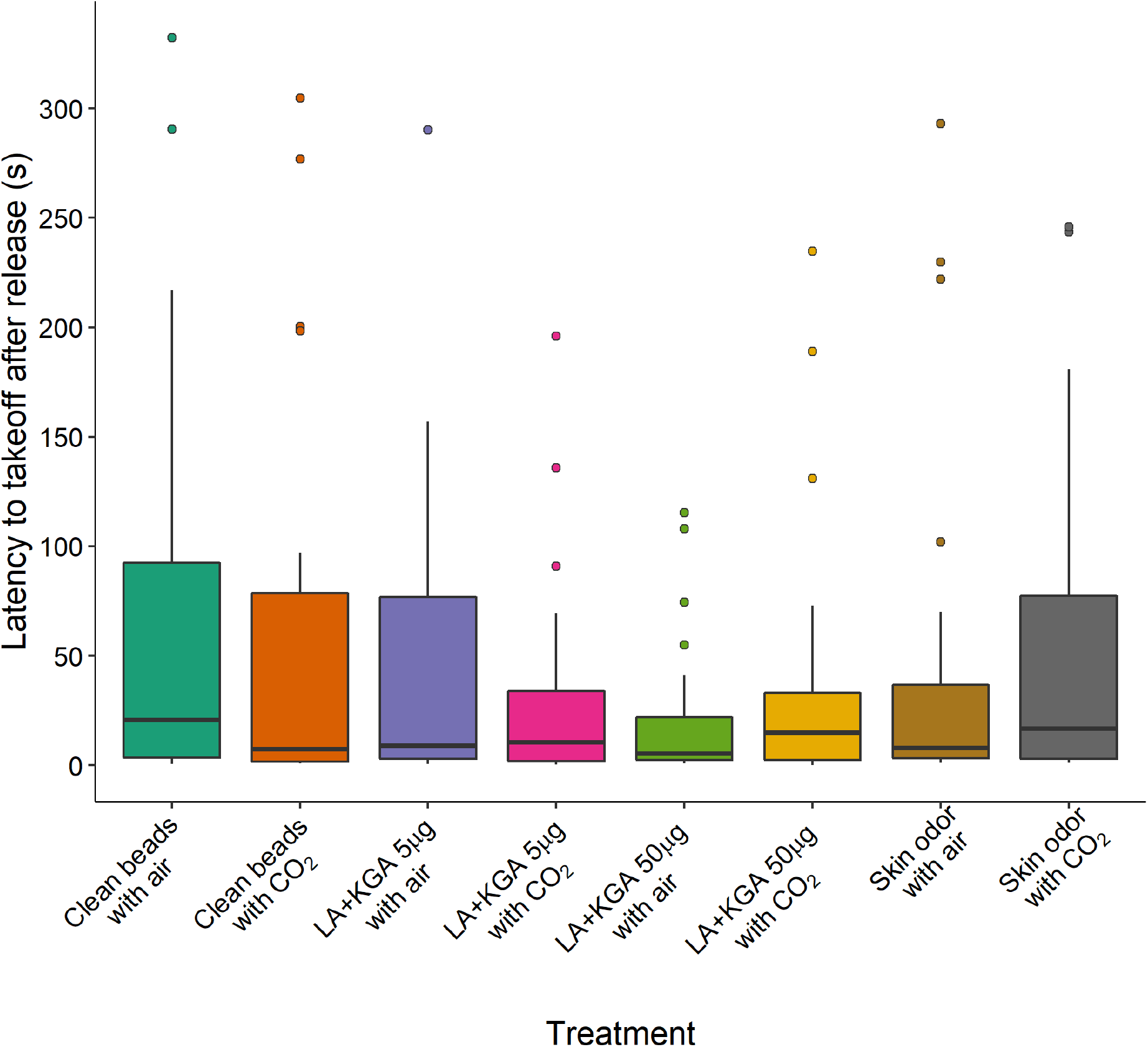
Latencies in *Aedes aegypti* of takeoff in seconds. While the high dose of the blend was significantly associated with a shorter latency (GLM, Est. = −0.3534, P = 0.017), the residuals of the GLM (gamma, link = log), despite a square root transform of the data, were somewhat divergent from the normal distribution (KS P = 0.022).

### Time From First Takeoff to First Landing

Figure 3 shows the time from takeoff to the first landing on the beads. Skin odor (Est. = −1.88, P < 0.001) and lactic and ketoglutaric acids 50 μg (Est. = −1.061, P = 0.026) reduced the time mosquitoes took to first reach and land on the beads. The presence of CO_2_ was not correlated with duration of flight from takeoff to first landing (P = 0.139).

**Fig. 3.**
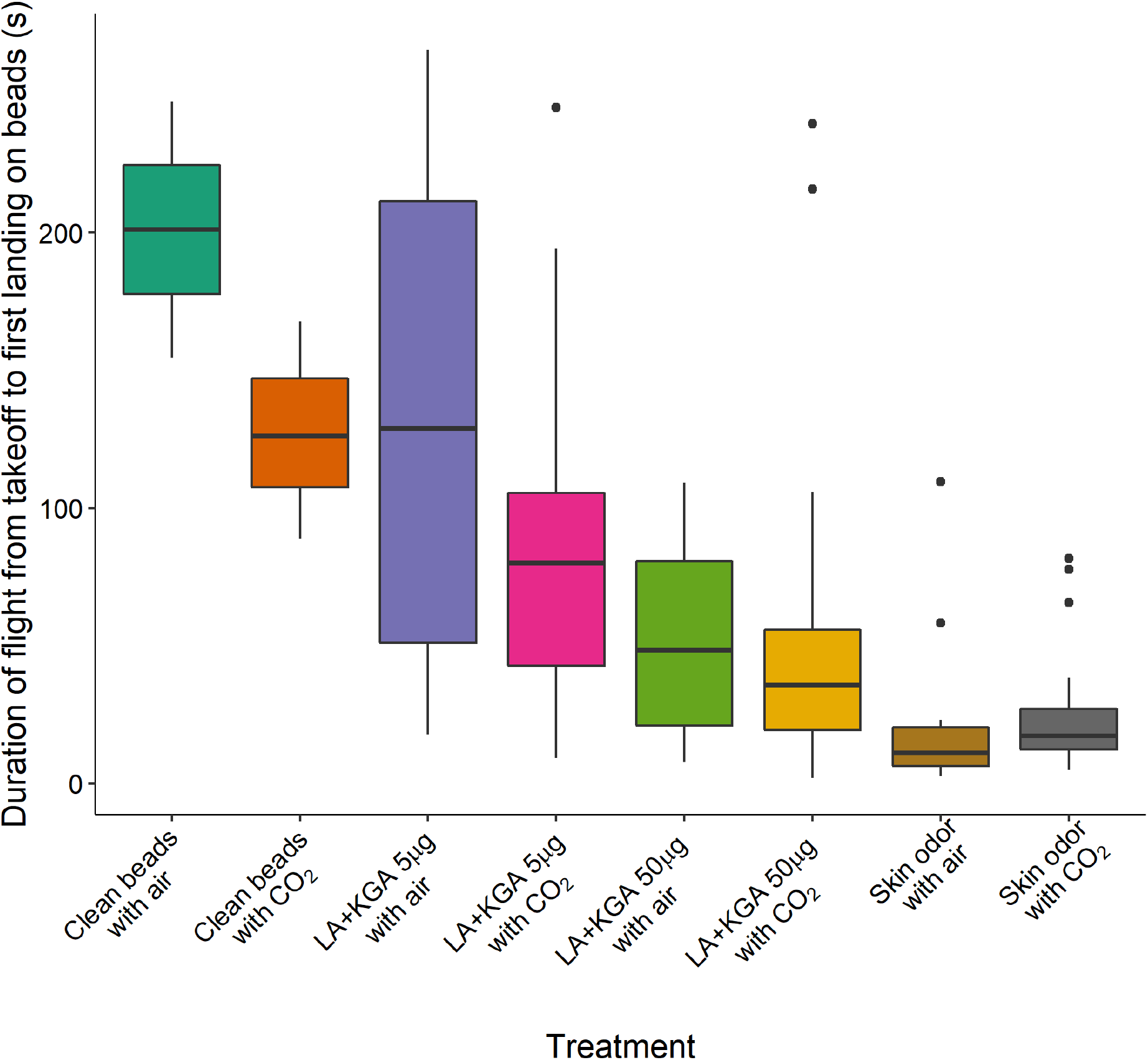
The durations of *Aedes aegypti* flight from takeoff to the first landing on beads in trials with at least one landing. The presence of skin odor (Est. = −1.88, P < 0.001) and the blend of 50 μg each of lactic and ketoglutaric acids (Est. = −1.061, P = 0.026) resulted in shorter flight times from takeoff to first landing. CO_2_ was not correlated with flight duration.

### Proportion of Trials with at Least One Landing

While skin odor with or without supplemental CO_2_ elicited the numerically highest proportion of trials with one or more landings on the odor-treated beads, it was not significantly different from the proportion of trials with lactic and ketoglutaric acids 50 μg in the presence or absence of CO_2_ that elicited one or more landings (Fisher Exact Test, adjusted P = 0.402) (Fig. 4).

**Fig. 4.**
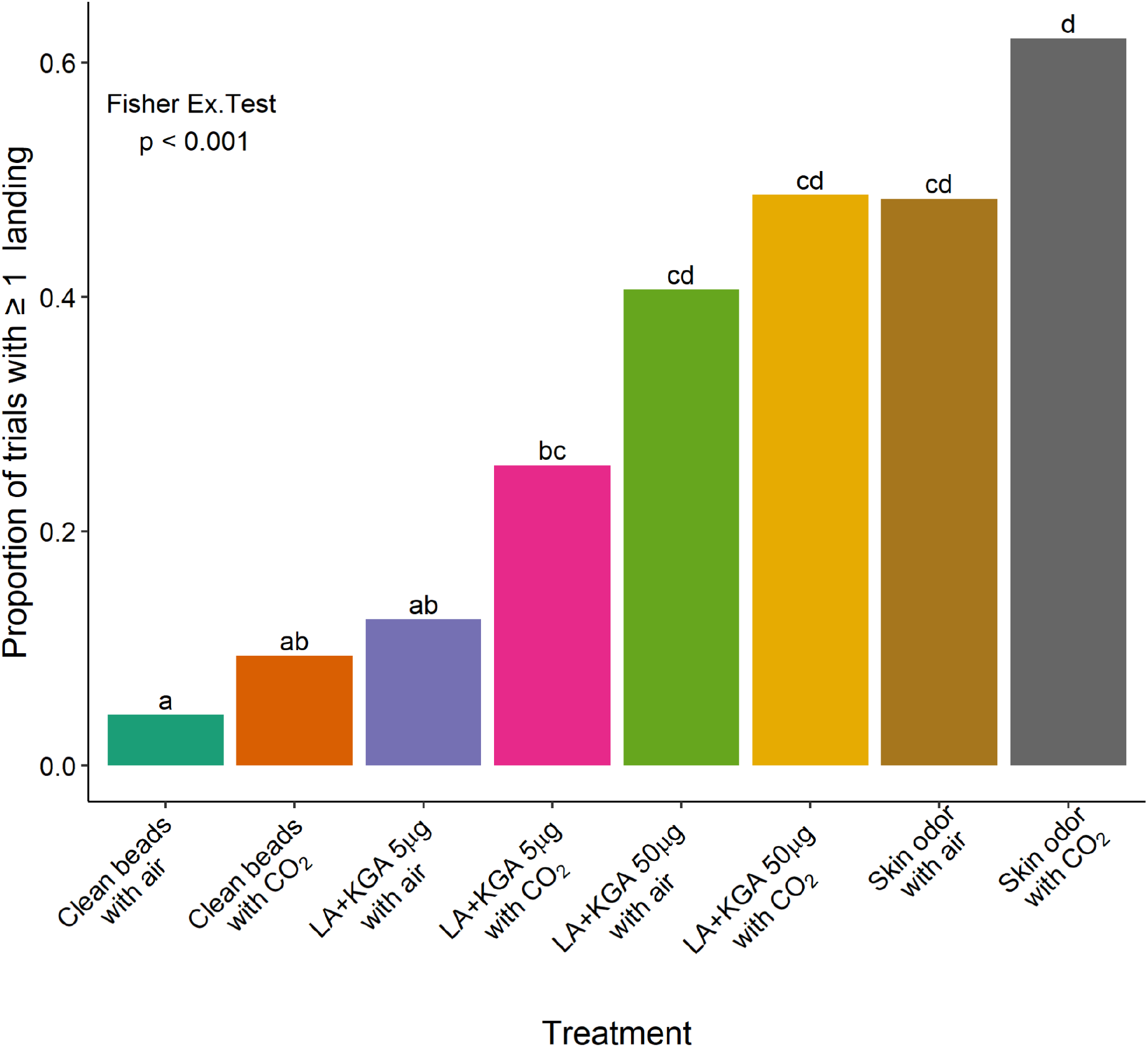
The proportion of trials in which *Aedes aegypti* landed more than once was tested with a Fisher Exact test followed by pairwise Fisher Exact tests; letters above columns show significant difference (P < 0.05). The GLMs, informed of the relationships among the treatments, found that: the proportion of all trials with ≥ 1 landing is correlated with: CO_2_ (Est. = 0.578, P = 0.046), skin odor (Est. = 2.88, P < 0.001), the blend of 5 μg each of lactic and ketoglutaric acids (Est. = 1.192, P = 0.029), and the blend of 50 μg of each (Est. = 2.434, P < 0.001).

Only the presence of skin odor (GLM, Est. = 2.88, P < 0.001) and both doses of the blend of lactic and ketoglutaric acids 50 μg (GLM, Est. = 2.434, P < 0.001) and 5 μg (GLM, Est. = 1.192, P = 0.029), were positively correlated with the probability of a mosquito landing at least once during a trial in the wind tunnel.

### Repeat Landings

Figure 5 shows the number of repeat landings, per trial by treatment, among trials with at least one landing. The presence of skin odor (Est. = 0.3238, P = 0.0199) and the high dose blend, 50 μg each, of lactic and ketoglutaric acids (Est. = 0.3244, P = 0.0311) were positively correlated with the landings by a single mosquito per trial (Fig. 5).

**Fig. 5.**
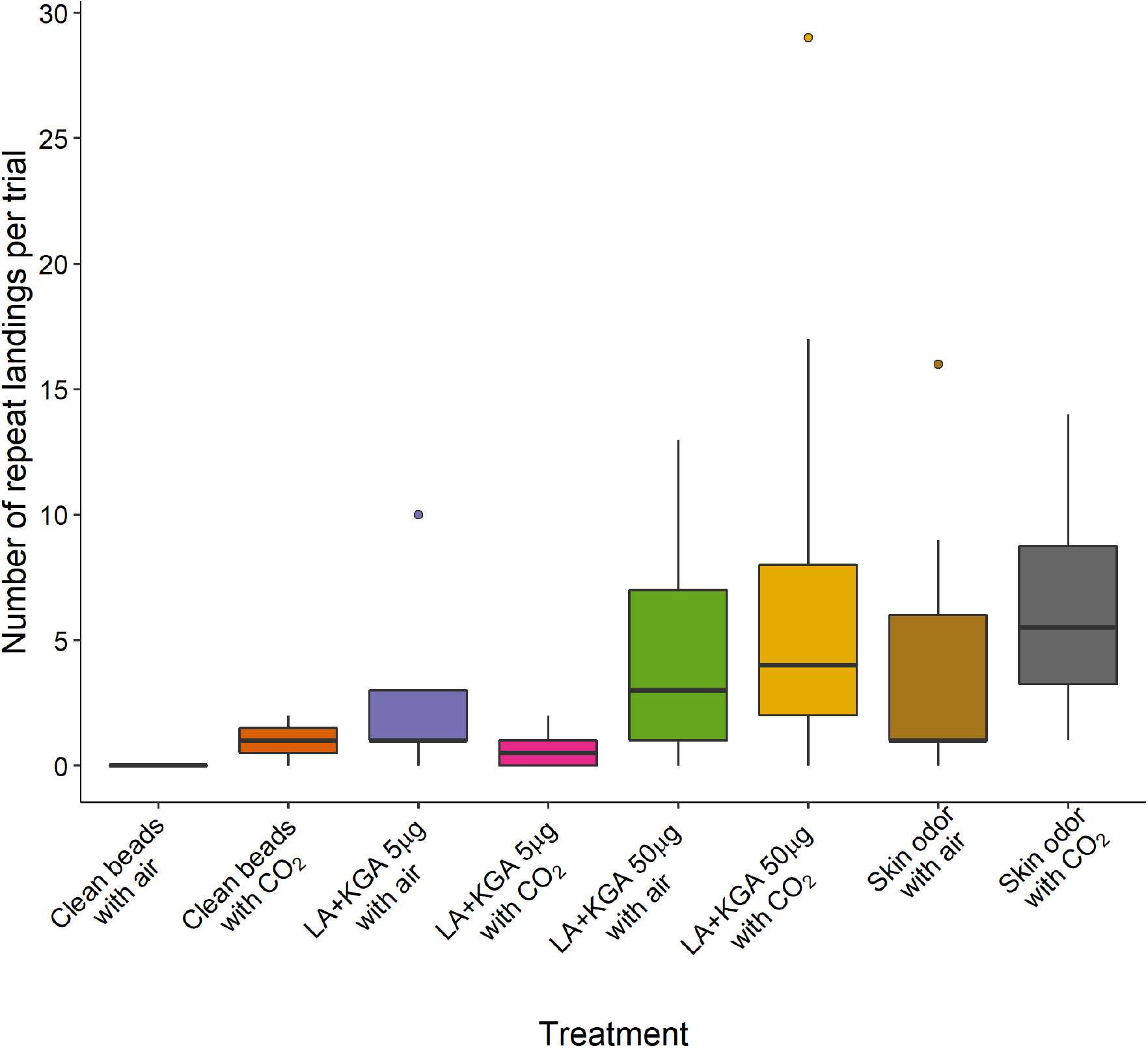
The number of repeat landings by female *Aedes aegypti* per trial among trials with at least one landing. Only skin odor (Est. = 0.3238, P = 0.0199) and 50 μg each of lactic and ketoglutaric acids (Est. = 0.3244, P = 0.0311) were correlated with the number of landings per trial.

### Total Landings on Lactic Acid Alone or the Lactic and Ketoglutaric Acid Blend

The number of landings on the 50 μg each blend of lactic and ketoglutaric acids (12.2 mean landings per trial, S.D. = 8.9) were significantly greater than those on lactic acid alone (5.4 mean landings per trial, S.D. = 6.5) (Kruskal-Wallace, P = 0.046) (Fig. 6). The time from takeoff to the first landing was significantly shorter between the two-component blend (mean = 57 seconds, S.D. = 49.4) and the lactic acid alone (mean = 115 seconds, S.D. = 99.3) (Kruskal-Wallace, P = 0.039).

**Fig. 6.**
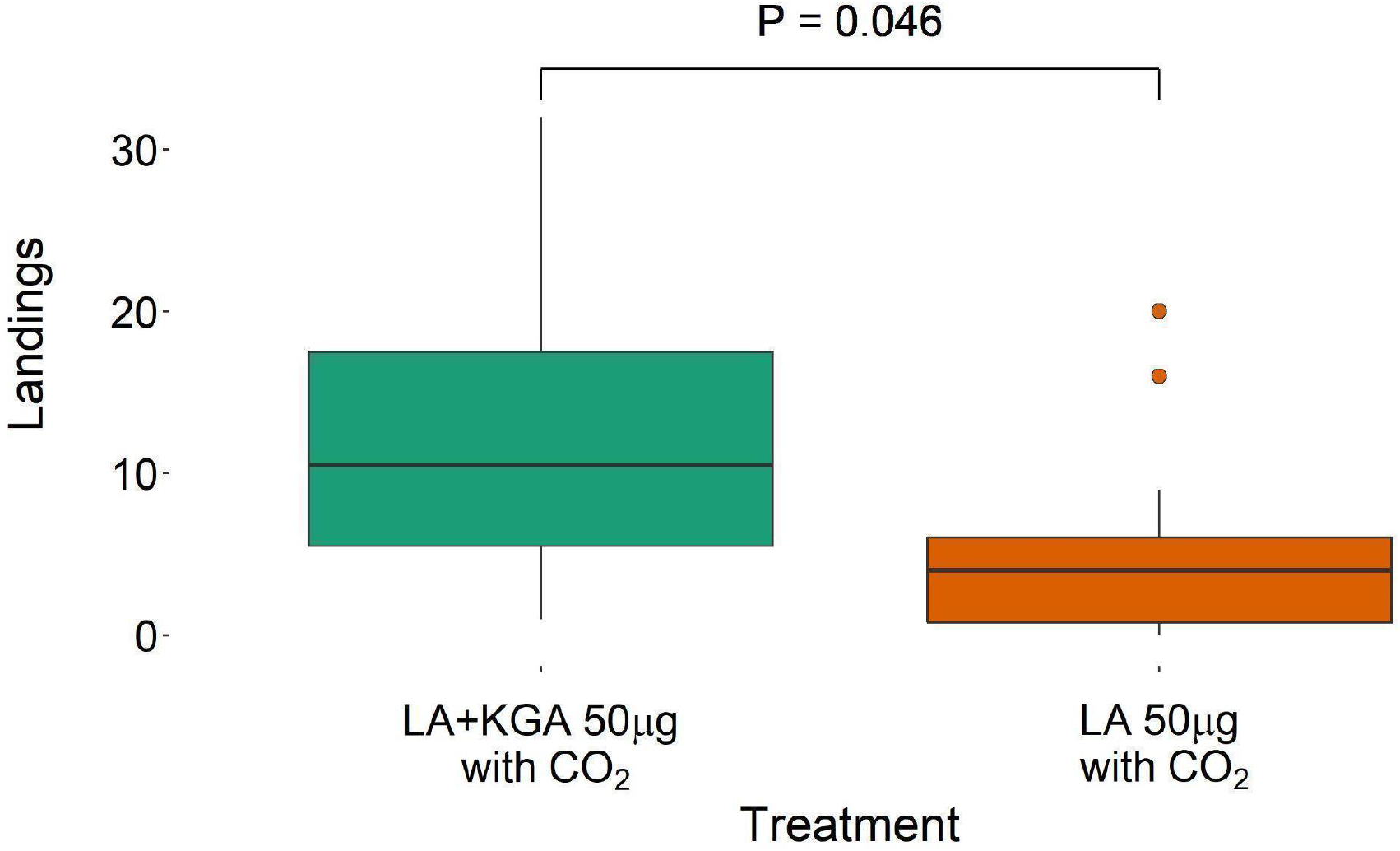
The total landings of *Aedes aegypti* by trial with the blend versus lactic acid alone. The number of landings on the two-component blend was significantly different than on lactic acid alone (Kruskal-Wallace, P = 0.046).

### Duration of Landing

Figure 7 provides the time within each trial that a single mosquito spent on the beads. The skin odor treatment (Est. = 1.626, P < 0.001), both doses of the blend of lactic and ketoglutaric acids, 5 μg (Est. = 0.979, P = 0.028), and 50 μg (Est. = 1.207, P = 0.004) were positively correlated with the duration of landing time of the mosquito (Fig. 7). The presence of CO_2_ was not correlated with duration of landing time. Additionally, mosquitoes were observed sticking their proboscises on the beads coated with 50 μg each of lactic and ketoglutaric acids, in the presence of CO_2_, in a manner resembling probing behavior.

**Fig. 7.**
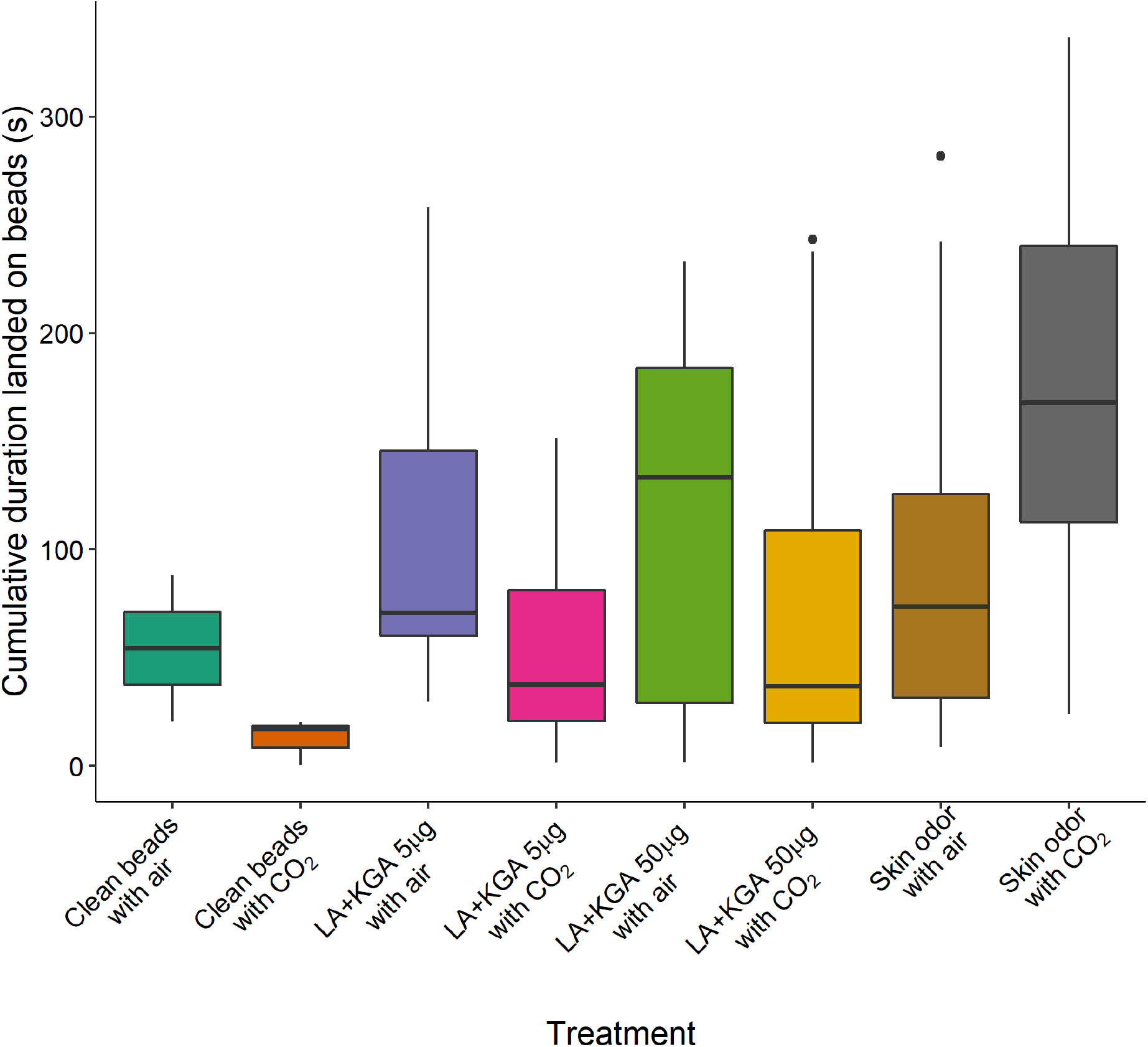
The cumulative duration of times female *Aedes aegypti* spent on beads, in trials with ≥ 1 landing. Skin odor (Est. = 1.626, P < 0.001), 5 μg each of lactic and ketoglutaric acids (Est. = 0.979, P = 0.028), and the blend of 50 μg of each (Est. = 1.207, P = 0.004) were correlated with the duration mosquitoes remained landed. CO_2_ was not correlated with duration landed.

### Analysis of Flight Tracks from Takeoff to First Landing

Among the treatments there were no differences in the mean distance of insects from the beads during flight in the wind tunnel (χ^2^ = 298, df = 298, P = 0.49). Selected flight tracks are shown in Fig. 8. Across all treatments, the mean (± SE) distance (mm) from the center of the beads during a mosquito flight was 214.15 mm (± 5.47). This mean distance from the beads was negatively correlated with whether an individual landed on the beads (ρ = −0.36, P < 0.001).

**Fig. 8.**
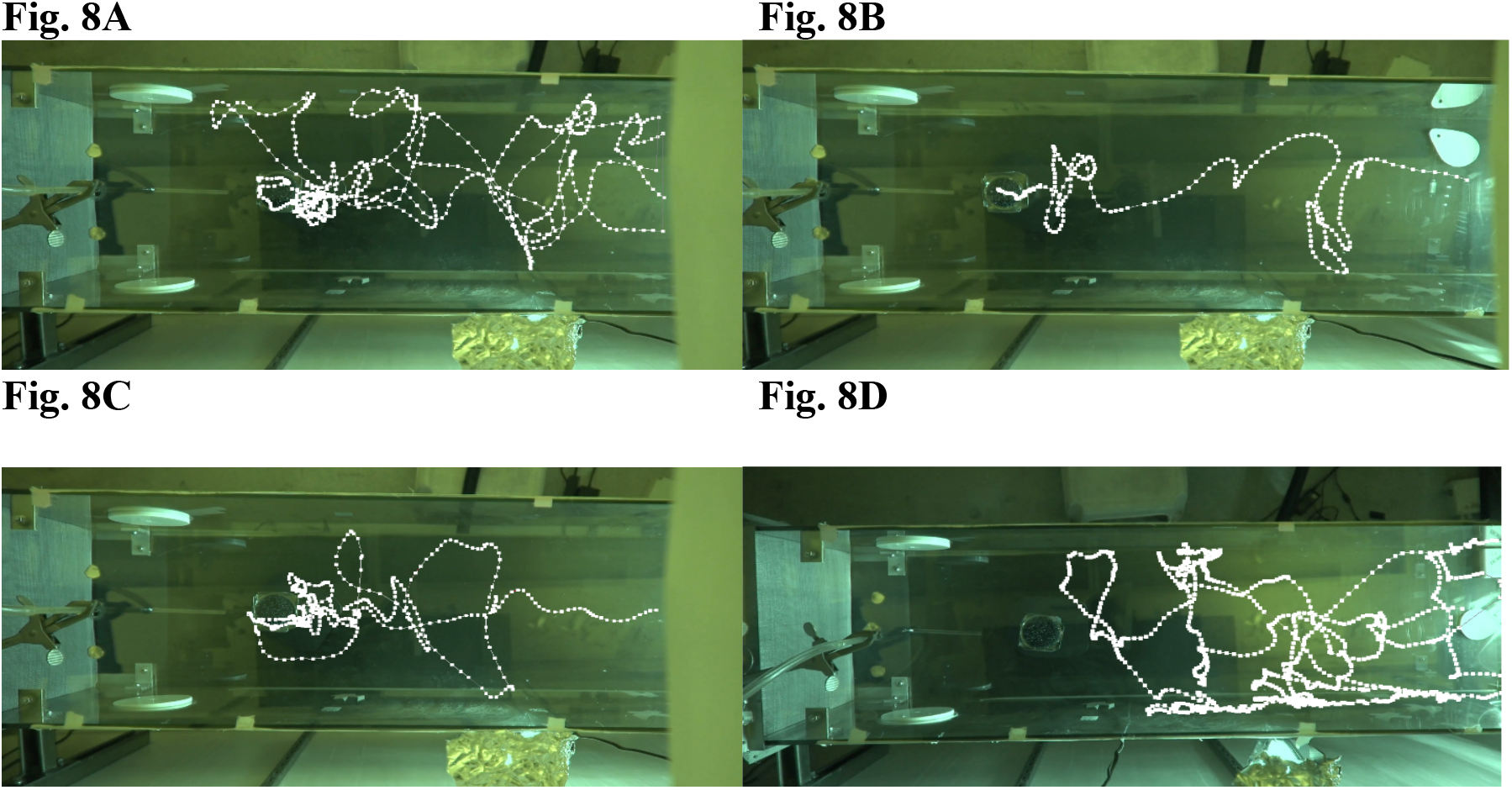
Top view of selected flight tracks of female *Aedes aegypti*, provided with CO_2_, responding to A) 50 μg of lactic and ketoglutaric acids; B) 5 μg of lactic and ketoglutaric acids; C) Skin odor; D) Clean beads. The mosquito did not land in the track shown in 8D. Airflow was left to right, and the mosquitoes were released on the right.

There were no differences in the mean flight velocity (mm/s) of insects during flight in the wind tunnel among treatments (χ^2^ = 298, df = 298, P = 0.49), nor did velocity of an individual significantly correlate with whether that individual landed on the beads or not (ρ = −0.12, P = 0.05). Across all treatments, the mean (± SE) velocity (mm/s) of mosquito flight in the wind tunnel was 193.15 mm (± 3.67).

There were no differences in the proportion of time an insect spent heading (± 20°) toward the center of the beads during flight in the wind tunnel among treatments (χ^2^ = 289.36, df = 288, P = 0.47), nor did this proportion of time significantly correlate with whether that individual landed on the beads or not (ρ = 0.01, P = 0.83). Across all treatments, the mean (± SE) proportion of the time a flying mosquito spent heading towards the center of the beads (± 20 °) was 0.16 (± 0.0039).

The E_max_ (track straightness; 1 = completely straight track) of insect tracks during flight in the wind tunnel did not differ among treatments (χ^2^ = 298, df = 298, P = 0.49), nor did the E_max_ of an individual significantly correlate with whether that individual landed on the beads or not (ρ = 0.05, P = 0.44). Across all treatments the mean (± SE) E_max_ of mosquito flight in the wind tunnel was 0.35 (± 0.01).

## Discussion

The two-component blend of lactic acid and ketoglutaric acid developed by Bello and Cardé (2022) elicited upwind flight and landing. The proportion of *Ae. aegypti* that landed at least once on an upwind odor source of the “high dose” (50 μg of each) of the blend was similar to the proportion that landed on human a source of man odor. Unsurprisingly, the presence of CO_2_ also increased the proportion of landings, given the known role of CO_2_ in sensitizing mosquitoes to human skin odors (Dekker et al. 2005) and eliciting upwind flight (Kennedy 1940). Individual humans vary in their intrinsic attractiveness to mosquitoes, and some of this variation is likely attributable to the quantitative differences among individuals in their emission of lactic acid and ketoglutaric acid (Thurmon and Ottenstein 1952; Delgado-Povedano et al. 2020).

Skin odor and the higher dose of the blend of lactic acid and ketoglutaric acid (50 μg), induced rapid orientation and landing on the beads and comparable numbers of repeat landings per individual mosquito. This suggests that this blend elicited the same persistence in mosquito landing behavior as human skin odor. The similarity between the behavioral activity elicited by the high dose and skin odor was further demonstrated in that the blend and skin odor were both correlated with the cumulative duration of landing on the beads. The number of repeat landings, their durations, or latency of first landing were not correlated with the presence of CO_2_. This is unsurprising, because in nature a mosquito landing on a skin odor source other than skin on the face would have likely exited the CO_2_ plume (Dekker and Takken 1998). Because a long-duration landing reduces the available time for further landings, the numbers and durations of landings were inversely correlated in all trials.

The high-dose blend of lactic acid and ketoglutaric acid (50 μg) elicited more landings than lactic acid alone. We found that 50 μg rather than 5 μg of each component of the blend elicited *Ae. aegypti* behaviors akin to skin odor in a wind tunnel. The 5 μg dose was sufficient to induce numerous landings the cage assay which has little ambient air movement (Bello and Cardé, 2022). The need for a higher dose to evoke upwind source finding in a wind tunnel is consistent with the odor mixture being diluted by turbulent diffusion of the wind-borne plume as it is carried downwind.

Carbon dioxide was positively correlated with the probability of takeoff. It was not correlated with the proportion of trials with a landing within the subset of trials with mosquito take off. Our findings support that CO_2_ elicits takeoff in *Ae. aegypti* females but does not act as a landing cue. This is consistent with the sequential-distance model of mosquito host seeking (Gillies 1980; Cardé and Gibson 2010; van Breugel et al. 2015; Cardé 2015). In this paradigm a host seeking mosquito takes off and flies upwind in a plume of CO_2_ before encountering other cues. The mosquito is then able to detect visual cues, host odors other than CO_2_, and finally heat from the host. There are physical limits to the distance at which visual and heat cues should be detectable to mosquitoes (Kahn et al. 1966; Muir et al. 1992).

Ketoglutaric acid is a component of the citric acid cycle (Wishart et al. 2018). It is found in fresh and dry sweat (Delgado-Povedano et al. 2020). It is not known how much ketoglutaric acid volatilizes from human skin. Lactic acid is released from human apocrine glands at a rate exceeding that of many non-human animals (Thurmon and Ottenstein 1952). Incubated sweat contains less lactic acid than fresh sweat, suggesting that most is produced endogenously (Braks and Takken 1999).

Lactic acid has been a controversial candidate as a mosquito attractant (Acree et al. 1968; Smith et al. 1970); Steib et al. (2001) added lactic acid to human and non-human animal odor. The addition of lactic acid resulted in the non-human animals’ odor drawing as many *Ae. aegypti* to its arm of the Y-tube as human odor. However, lactic acid alone attracted only 19 % of the mosquitoes tested. Calf and goat odors with added lactic acid attracted 70 % of the *Ae. aegypti* tested.

Ketoglutaric acid may be one of the compounds Steib et al. (2001) and Geier et al. (2002) demonstrated existed but did not isolate. Our results corroborate those of Bello and Cardé (2022), that lactic acid is a necessary component but insufficient alone to elicit a rate of mosquito landing equal to that of a blend of human odor compounds.

The time to takeoff was surprisingly similar across treatments. Even across those with and without CO_2_ the differences were not as large as we would have expected. We suspect that the release, while conducted with care, may have mechanically disturbed the mosquitoes enough to influence takeoff. A short latency to takeoff, however, may be an intrinsic character of *Ae. aegypti*. Cilek et al. (2004) described this mosquito as opportunistic or exhibiting “aggressive biting” and found that time to biting after landing averaged 9.8 ± 0.3 s (as opposed to *Culex quinquefasciatus*, which averaged 41.0 ± 1.1 s). The test of five mosquitoes at a time with the two-component blend and lactic acid alone was intended to determine if the blend was better at eliciting landing than lactic acid alone. The blend elicited more landings and those mosquitoes landed more quickly than to lactic acid alone.

The lack of statistical differences in flight racks among treatments was unexpected, as there are large differences in the landing propensities among treatments. It may be that such tracks are inherently “messy,” and they do not differ in flight characteristics in our assay or in our method of analysis.

Skin odor elicited longer landing durations than either dose of the synthetic lure. This may be in part due to the different suite of cues available after the mosquitoes contacted the beads. Mosquitoes have express gustatory receptors on their tarsi (Sparks et al. 2013). Along with chemoreceptors on the labellum (Saveer et al. 2018), receptors on the tarsi mean that after landing a mosquito may bring chemoreceptors into direct contact with host cues. These may include non-volatile chemicals such as amino acids. Further research, possibly with a cage-landing assay, will be needed to identify possible post-landing cues.

Current mosquito traps and lures, including the widely deployed BG Sentinel, have low trapping efficiency (actually captured after being lured to the trap’s vicinity) and require a fan (Amos et al. 2020 a,b; Amos and Cardé 2022). By using compounds that elicit landing, it might be feasible to lure the mosquitoes directly into traps. This would boost trap efficiency and perhaps allow the development of traps without fans. We counted the number of repeat landings. If a lure in a trap elicits repeated landing attempt, it would provide multiple opportunities for capture.

Many other compounds also are reported to be attractive to *Ae aegypti* mosquitoes (Coutinho-Abreu et al. 2021; Dormont et al. 2021) and should be evaluated to determine if any of these add to the attractiveness of the blend of lactic and ketoglutaric acids. Among these are hexanoic acid, a known attractant of *Ae. aegypti* (Carlson et al. 1973; Williams et al. 2006; Owino et al. 2015), and ammonia (Steib et al. 2001). While known as an attractant of *Anopheles* rather than *Aedes* mosquitoes, 2-oxopentanoic acid, as well as straight-chain carboxylic acids of various lengths (Healy and Copeland 2000; Healy et al. 2002) should be evaluated, although carboxylic acids and short-chain aldehydes can be repellent to *Ae. aegypti* (Logan et al. 2008; Owino et al. 2015). To avoid specious interpretations of which odors mediate natural attraction to human hosts, it will be important to release odors at rates and ratios closely mimicking those that are naturally emitted (Cardé 2022).

## Conclusion

A blend of lactic and ketoglutaric acids discovered by Bello and Cardé (2022) elicits upwind flight and landing of *Ae. aegypti* in a wind tunnel, with and without supplemental CO_2_. The effectiveness of this blend without supplemental CO_2_ makes this combination a candidate for use in mosquito traps. Ketoglutaric acid may be one of the compounds present in animal odors that when supplemented with lactic acid are highly attractive to anthropophilic mosquitoes.

## Acknowledgments

We are grateful to Dr. ES Lacey for experimental advice and assistance with rearing. Drs. AC Gerry and JG Miller provided useful comments on an early version of this manuscript. Dr. E Sarro provided statistical guidance.

## Data Availability

GitHub link to follow.

## Code Availability

GitHub link to follow.

## Ethics Approval

Not applicable.

## Consent to Participate

Not applicable.

## Consent for Publication

Not applicable.

## Open Access

## Authors’ Contributions

BDS and RTC designed the experiments, BDS carried out these trials and analyzed our findings. BAA analyzed the flight racks. JB aided in selection of treatments. BDS and RTC wrote the paper.

